# Modulation of Voltage-Gated Sodium Channel (VGSC) Activity in Human Dorsal Root Ganglion (DRG) Neurons by Herpesvirus Quiescent Infection

**DOI:** 10.1101/714691

**Authors:** Qiaojuan Zhang, Feng Chen, Miguel Martin-Caraballo, Shaochung V. Hsia

## Abstract

The molecular mechanisms of pain associated with alphaherpesvirus latency are not clear. We hypothesize that the voltage-gated sodium channels (VGSC) on dorsal root ganglion (DRG) neurons controlling electrical impulses may have abnormal activity during viral latent infection and reactivation. We used HSV-1 to infect human DRG-derived neuronal cell line HD10.6 to study viral latency establishment, maintenance and reactivation as well as changes of VGSC functional expression. Differentiated cells exhibited robust tetrodotoxin (TTX) sensitive sodium currents and acute infection significantly reduced the VGSC functional expression within 24 hours, and completely abolished the VGSC activity within three days lytic infection. A quiescent state of infection mimicking latency can be achieved by HSV infection in the presence of acyclovir (ACV) for 7 days followed by 5 days of ACV washout and the viruses can remain dormant for another three weeks. It was noted that during HSV-1 latency establishment, the loss of VGSC activity caused by HSV-1 infection could not be blocked by ACV treatment within 3 days infection. However, neurons with continued treatment of ACV for another 4 days showed a gradual recovery of VGSC functional expression. Furthermore, the latent neurons exhibited higher VGSC activity in comparison to controls. The overall regulation of VGSC by HSV-1 during its quiescent infection was proved by increased transcription and possible translation of Nav1.7. Together these observations demonstrated a very complex pattern of electrophysiological changes during HSV infection of DRG neurons, which may have implication for understanding the mechanisms of virus-mediated pain linked to latency and reactivation.

**Importance:** The reactivation of the herpesvirus from ganglionic neurons may cause cranial nerve disorders and unbearable pain. It is unclear why normal stimuli would trigger enhanced pain sensation in these patients. The current work meticulously studies the functional expression profile changes of VGSC during the process of latency establishment using an *in vitro* model. Our results indicated that VGSC activity was eliminated upon infection but steadily recovered during the latency establishment and the latent neurons exhibited even higher VGSC activity. This finding advances our knowledge of how ganglion neurons generated uncharacteristic electrical impulses due to abnormal VGSC functional expression influenced by latent virus.

## Introduction

HSV-1 is a neurotropic pathogen associated with excruciating orofacial sensation and other chronic neuropathic pains (1–9). After the acute infection, the virus may establish a lifelong latency in the neurons of trigeminal ganglion as well as other sensory and autonomic ganglia, including the dorsal root, geniculate, etc. through innervating sensory nerves (10). Reactivation may occur in response to various stress signals and trauma (11). The resulting pain and other complications cause detrimental impact on patients’ lives, causing considerable economic and health burden.

A variety of ion-channels and receptors are expressed in ganglion neurons such as transient receptor potential (TRP) channels, voltage-gated sodium channels, acid-sensing ion channels, ATP-sensitive receptors, potassium channels, voltage-gated calcium channels, and glutamate receptors (12). Nerve injury in dorsal root ganglion (DRG) neurons has been shown to affect their excitability and firing regardless injured or not (13, 14). It is known that patients with HSV-1 complication may experience hypoalgesia, dysesthesia, tingling, and formication but the underlying molecular mechanisms are unclear (8, 15, 16). Voltage-gated sodium channels (VGSCs) play a role in the generation and propagation of action potentials needed for the transmission and processing of pain signals (12). Changes in the molecular and/or functional expression of VGSCs during HSV-1 infections can potentially regulate sensory pain transmission by altering electrical excitability and the generation of electrical signals in pain-transmitting neurons (17). In addition, reduced electrical excitability has been shown to increase viral replication in HSV-1 infected neurons (18). These observations suggested that HSV-1 infection reduced neuronal excitability during early infection in order to enhance viral replication and further spread.

VGSCs consist of a multimeric complex containing a large α pore-forming subunit and several auxiliary subunits. In mammals, the main pore forming subunit is generated by the product of 10 genes (19). Of all these subunits, Nav1.1, 1.6, 1.7, 1.8, and 1.9 were distributed in DRG and TG (19) and Nav1.7, 1.8, and 1.9 were found to participate in mediating chronic pain in DRG (20). Disruption of Na^+^ channel expression has been associated with altered excitability in pain-transmitting neurons (21). Furthermore, inhibition of Na^+^ channels can reverse mechanical allodynia and thermal hyperalgesia in a model of herpes virus-induced post herpetic pain (22).

Our hypothesis is that HSV-1 quiescent infections can modulate the functional expression of Na^+^ channels and alter neuronal excitability, thus participating in the establishment of latency and regulation of pain. Changes in VGSC activity may underlie the sensory abnormalities described in patients suffering orofacial chronic pain associated with HSV-1 infection.

## Results

### Differentiated HD10.6 cells supported the establishment of HSV-1 quiescent infection and increased viral replication can be observed by TSA treatment

To test the capability of HD10.6 cells in supporting quiescent infection of HSV-1 strain McKrae/GFP, differentiated cells were infected at moi of 1 in the absence or presence of ACV followed by fluorescent microscopy to monitor the progress of viral infection. The results showed that without ACV HD10.6 cells were well-infected by McKrae/GFP and replicated efficiently at 3 dpi (compare Fig. 1A and B). In the presence of ACV, fluorescent microscopy indicated that approximately 60% of the cells were infected at 3 dpi but replication was significantly decreased as evidenced by the weak green fluorescence (Fig. 1C). The green fluorescence continued to decrease at 7 dpi (Fig. 1D) and was barely detected at 12 dpi after 5 days of ACV washout (Fig. 1E). This quiescent state of infection, found in approximately all of the infected cells, can be maintained for another two weeks without evidence of viral replication (data not shown). These quiescently infected cells, nonetheless, displayed significantly increased green fluorescence after adding the HDAC inhibitor TSA for 2 days (Fig. 1F). These findings suggested that differentiated HD10.6 cells were able to support HSV-1 infection in a dormant state mimicking latency and maintain the capacity to be rebooted to replicate similar to reactivation.

**Fig. 1.**
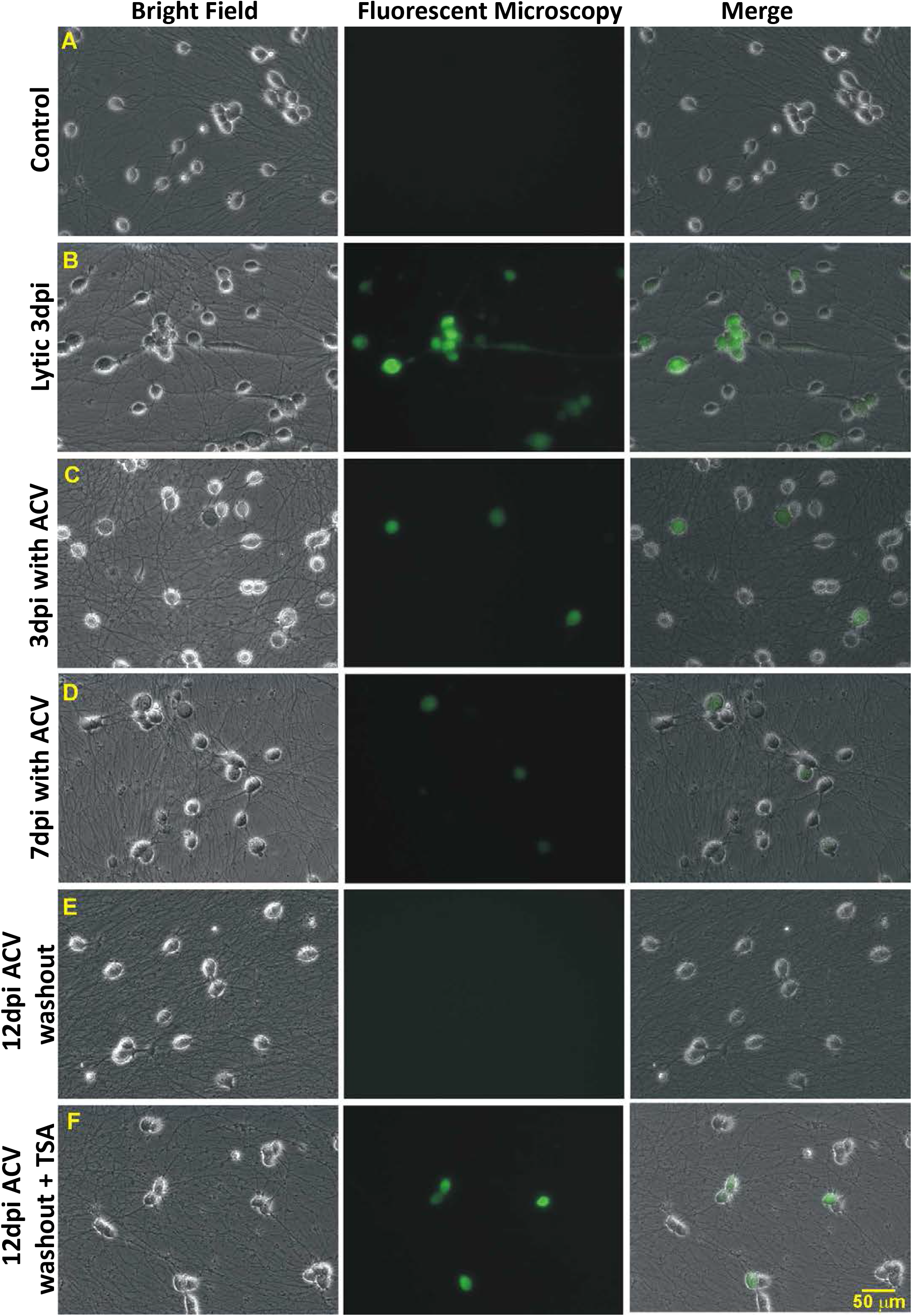
Establishing HSV-1 quiescent state of infection with ACV and triggering replication by TSA. Differentiated HD10.6 cells were subjected to HSV-1 infection followed by fluorescent microscopy examining the progress of infection. A) No infection control; B) Lytic infection, virus replicated and green fluorescence can be clearly seen at 3dpi; C) Infection with ACV at 3dpi, cells with green fluorescence can be observed with weaker intensity comparing to lytic infection; D) Infection with ACV at 7dpi, green fluorescent cells can be barely detected and the number decreased; E) Infection at 12dpi with 5d ACV washout, fluorescent microscopy can hardly detect the green fluorescent cells, suggesting a quiescent infection mimicking latency has been established and maintained. F) Infection at 12dpi with 3d ACV washout followed by 2d TSA treatment, fluorescent microscopy recorded more cells with stronger green fluorescence, suggesting the dormant state of infection has been disrupted with increased viral gene expression and replication, similar to reactivation observed in animal models.

### TSA increased viral gene expression and replication but no virus release

The effect of TSA was further analyzed by qRTPCR to investigate changes in viral gene expression. The results showed that HSV-1 TK and GFP under the control of CMV promoter were increased 3.47-fold and 124-fold by TSA, respectively, following reactivation (Fig. 2A and B). The PI3K inhibitor LY294002, which was used to induce HSV-1 reactivation (23–25), did not appear to increase the viral gene expression, probably due to the fact that neurotrophins such as NGF and GDNF were not removed (Fig. 2A and B). The release of infectious viral particles was tested by collecting media of infected HD10.6 cultures, followed by plaque assays. Neither the TSA nor the LY294002 treatment was able to trigger the release of viral particles (data not shown). The same experiment was performed using the infected HD10.6 cell lysate and the results indicated that TSA treatment produced a virus titer approximately 32 pfu/ml, while the latent cells and cells under LY294002 treatment generated no plaque (Fig. 2C). These results indicated that HDAC inhibitor TSA promoted viral gene expression and replication of the latent HD10.6 cells but the viral egress was impaired.

**Fig. 2.**
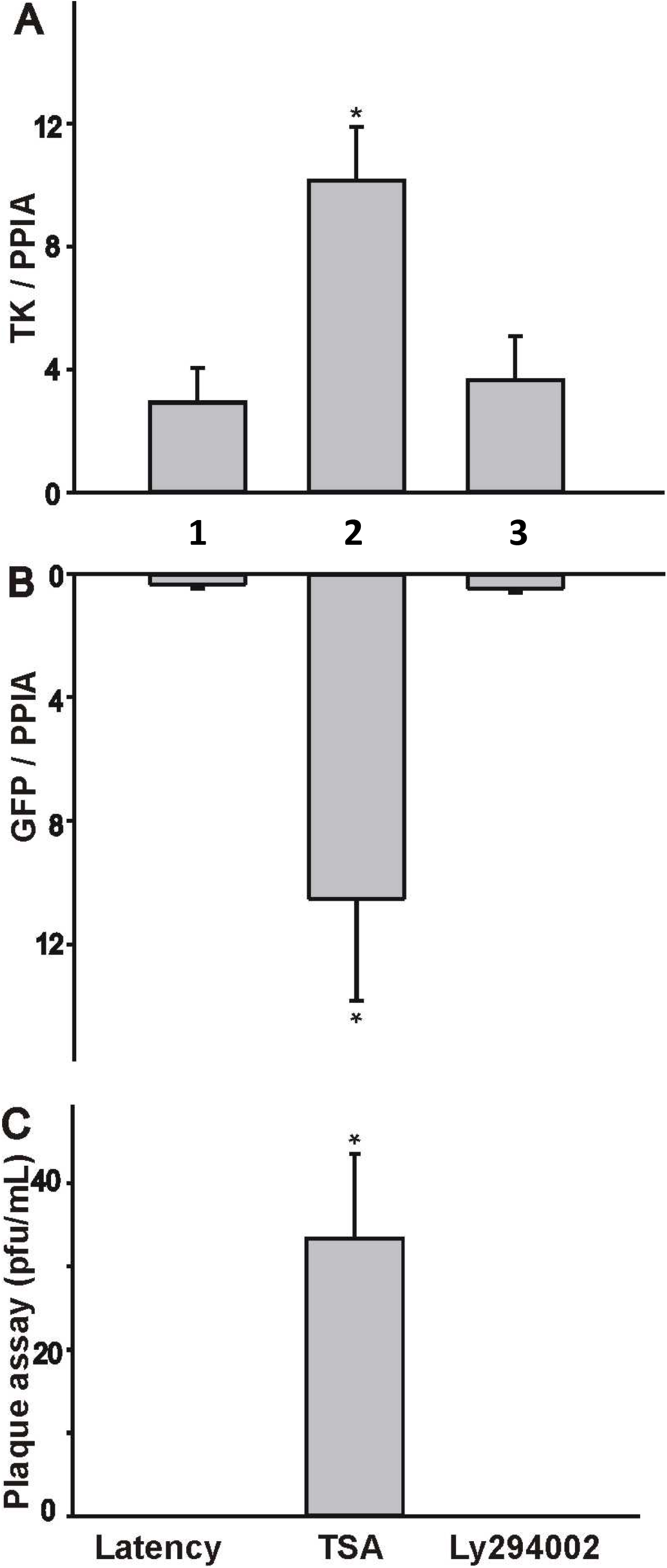
TSA induced viral gene expression and replication from the latent reservoir of HD10.6 cells. Viral latency and reactivation were assessed by qRTPCR and plaque assays, respectively. A) HSV-1 TK was analyzed by qRTPCR. Lane 1: Latent neurons without TSA nor LY294002; Lane 2: Latent neurons treated with TSA; Lane 3: Latent neurons treated with LY294002; B) GFP was measured by qRTPCR with the same arrangement. C) The plaque assays were performed using the cello lysates from infected HD10.6 cells. Lane 1: Lysate from latent neurons without treatment; Lane 2: Lysate from latent neurons treated with TSA; Lane 3: Lysate from latent neurons treated with LY294002. A symbol (*) denotes significant statistical difference with p value <0.05 vs latent group.

### HSV-1 gene expression decreased during the latency establishment and maintenance, but increased while reactivated

The protocol of latency establishment, maintenance, and reactivation was summarized in Fig. 3A. For latency establishment (LE), infected cultures were treated with 100 μM ACV for 7 days to initiate the formation of HSV-1 quiescent infection. For latency maintenance (LM), ACV was then removed to continue the dormant state of infection for 3 days. The reactivation was attempted by adding 1μM TSA for 2 days. Transcription of the viral genes ICP0, TK, and LAT in the absence and presence of ACV at different time points (3dpi, 7dpi, and 12dpi) was analyzed by qRTPCR. It was shown that, compared to lytic infection, ICP0 and TK decreased 50% and 75% at 3dpi in the presence of ACV, respectively (Fig. 3B). LAT, on the other hand, increased and accumulated to a comparable level at 3dpi without viral replication (Fig. 3B). All three viral transcripts, nonetheless, declined at 7dpi and 12dpi but TSA reversed the diminishing trend with significant increase (Fig. 3B). Collectively these results suggested that differentiated HD10.6 supported the establishment of HSV-1 quiescent infection thus mimicking the maintenance of latency. TSA treatment overturned the dormant state, increased the viral gene expression, and promoted replication. LAT did not accumulate when the virus established latency *in vitro* but was relatively abundant temporarily at 3dpi without viral replication.

**Fig. 3.**
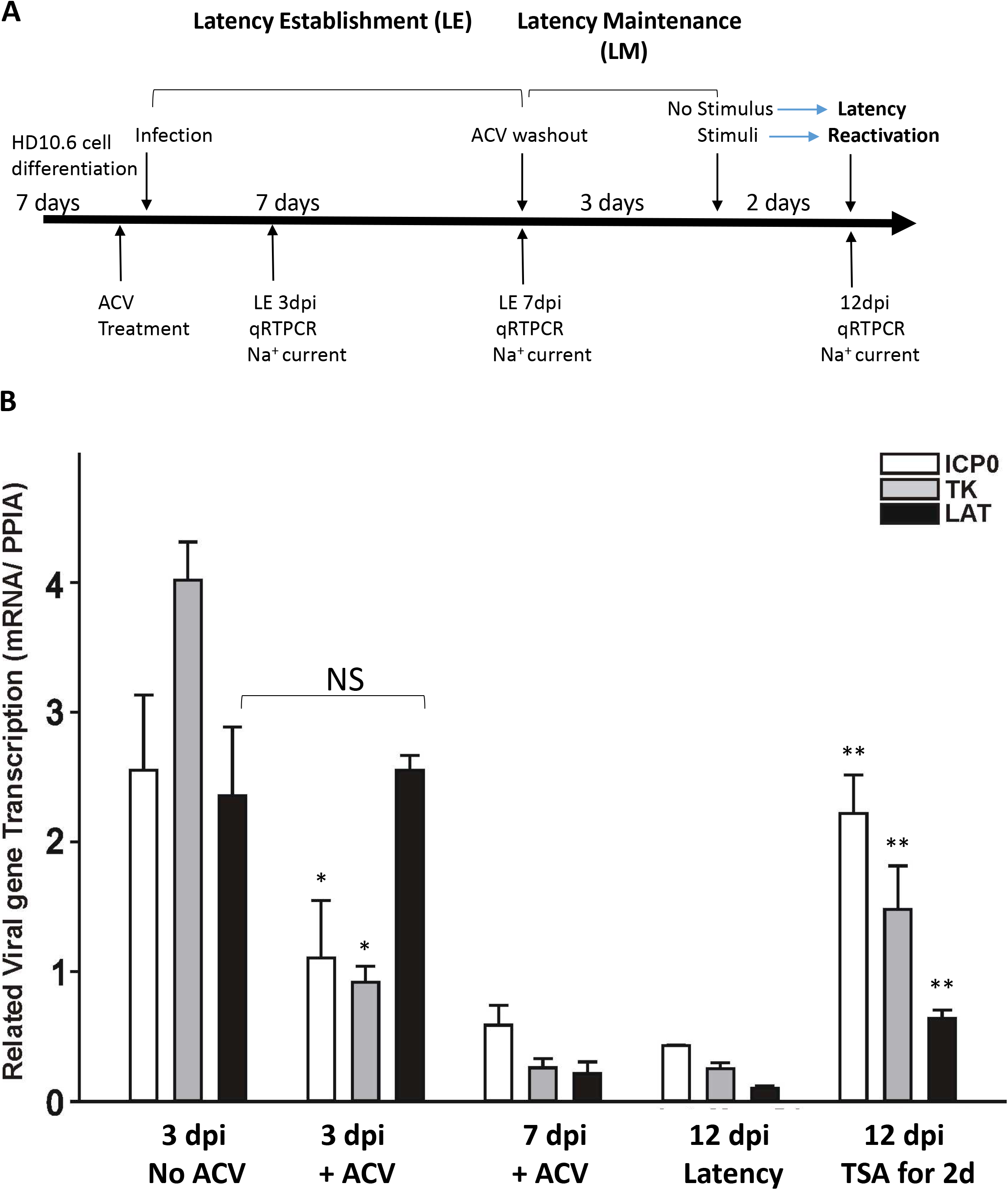
Transcription profile of three HSV-1 genes during the course of latency establishment and reactivation. A) Schematic of human DRG cell line derived neuronal system used to investigate HSV-1 latency establishment, maintenance and reactivation as well as the related time point for sodium current recording. B) The transcription of three viral genes (ICP0: white bar, TK: gray bar, and LAT: black bar) was assessed by qRTPCR at different time points (3dpi, 7dpi, and 12dpi) in the absence of viral replication, comparing to lytic infection at 3dpi and reactivation by TSA at 14dpi. A symbol (*) denotes significant statistical difference with *p* value <0.05 vs. correspondent gene in the 3dpi no ACV group. A double-asterisk (**) signifies significant statistical difference with *p* value <0.05 compared to respective gene in the 12dpi latency group. NS indicates not significant statistically.

### Na^+^ channel activity is abolished by HSV-1 infection but gradually recovered in latently-infected HD10.6

To test the impact of viral infection on sodium channel activity, differentiated cells were subjected to McKrae/GFP infection followed by electrophysiological recordings of Na^+^ currents from cells expressing green fluorescence as evidence of infection. VGSC activity was measured as current density, which was calculated by dividing the maximal current amplitude by cell capacitance (Fig. 4A). Differentiated control cells exhibited robust sodium currents and lytic infection at 3dpi eliminated the Na^+^ currents (Fig. 4B, Lane 3), similar to what we previously observed in ND7/23 cells (26). The effects of infection in the presence of ACV on the Na^+^ currents were assessed at 3dpi and 7dpi. It can be seen that Na^+^ currents were not affected by ACV in controls (Fig. 4B, Lane 2) but reduced at 3dpi with ACV (Fig. 4B, Lane 4). This is different from the previous report that ACV treatment can maintain VGSC activity after HSV-1 infection (27). Sodium current densities were recovered without significant difference comparing to control during the latency establishment (LE) at 7dpi (Fig. 4B, Lane 5). These results suggested that VGSC activity lost upon HSV-1 infection was rescued during LE.

**Fig. 4.**
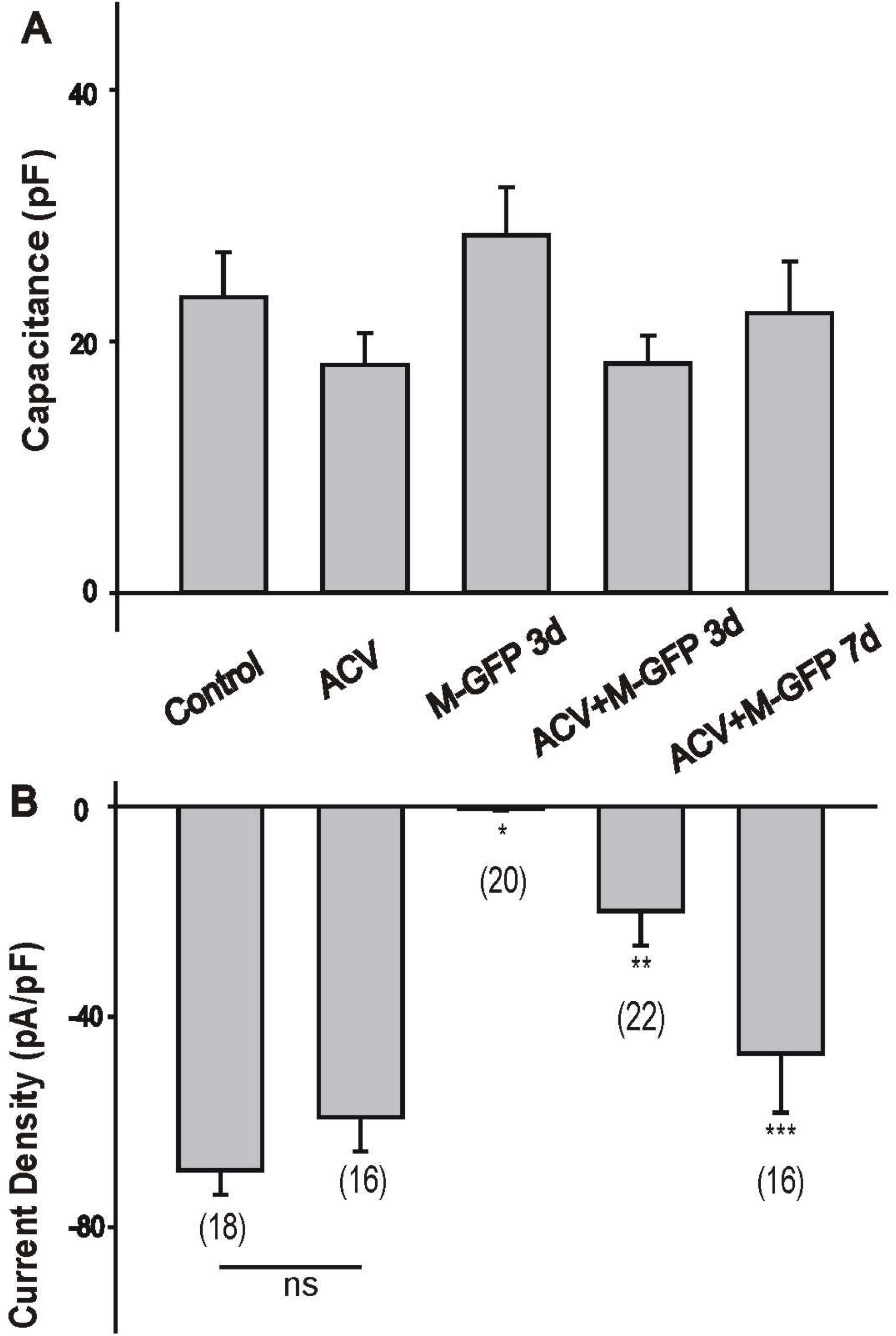
Sodium current recording during the HSV-1 latency establishment in HD10.6 cells. A) Comparison of cell capacitance in HD10.6 cells under differentiation conditions. No ACV treatment (Control), ACV treatment (ACV), HSV-1 lytic infection for 3 days (M-GFP 3d), latency establishment for 3 days (ACV+M-GFP 3d), latency establishment for 7 days(ACV+M-GFP 7d). B) The Mean Na^+^ current densities generated in HD10.6 cells under different conditions. Note the voltage-gated Na^+^ current density was calculated from the peak current amplitude dividing by the cell capacitance. The asterisk denotes p< 0.05 vs. control; the double asterisk denotes p < 0.05 vs. ACV group; the triple asterisk denotes p <0.05 vs. ACV+M-GFP 3d.

### HD10.6 cells exhibited robust TTX-sensitive Na^+^ currents

To assess the nature of HD10.6 VGSC, Na^+^ current was subsequently determined by TTX. It appeared by the analyses of Na^+^ current trace (Fig. 5A) and current-voltage (I–V) relationship (Fig. 5B) that TTX at 250 nM significantly reduced the Na^+^ current. Quantitative studies showed that approximately 88% of Na^+^ current was eliminated by TTX (Fig. 5C), in agreement with the previous finding that that HD10.6 cells primarily utilized TTX-sensitive VGSC for its physiological activity (28). The expression of two TTX-sensitive channels Nav1.3 and Nav1.7 was analyzed by qRTPCR and the results indicated that Nav1.7, but not Nav1.3, was steadily increased during LE (Fig. 5D). Together these studies suggested that lost VGSC activity due to viral infection was recovered because of Nav1.7 expression during LE.

**Fig. 5.**
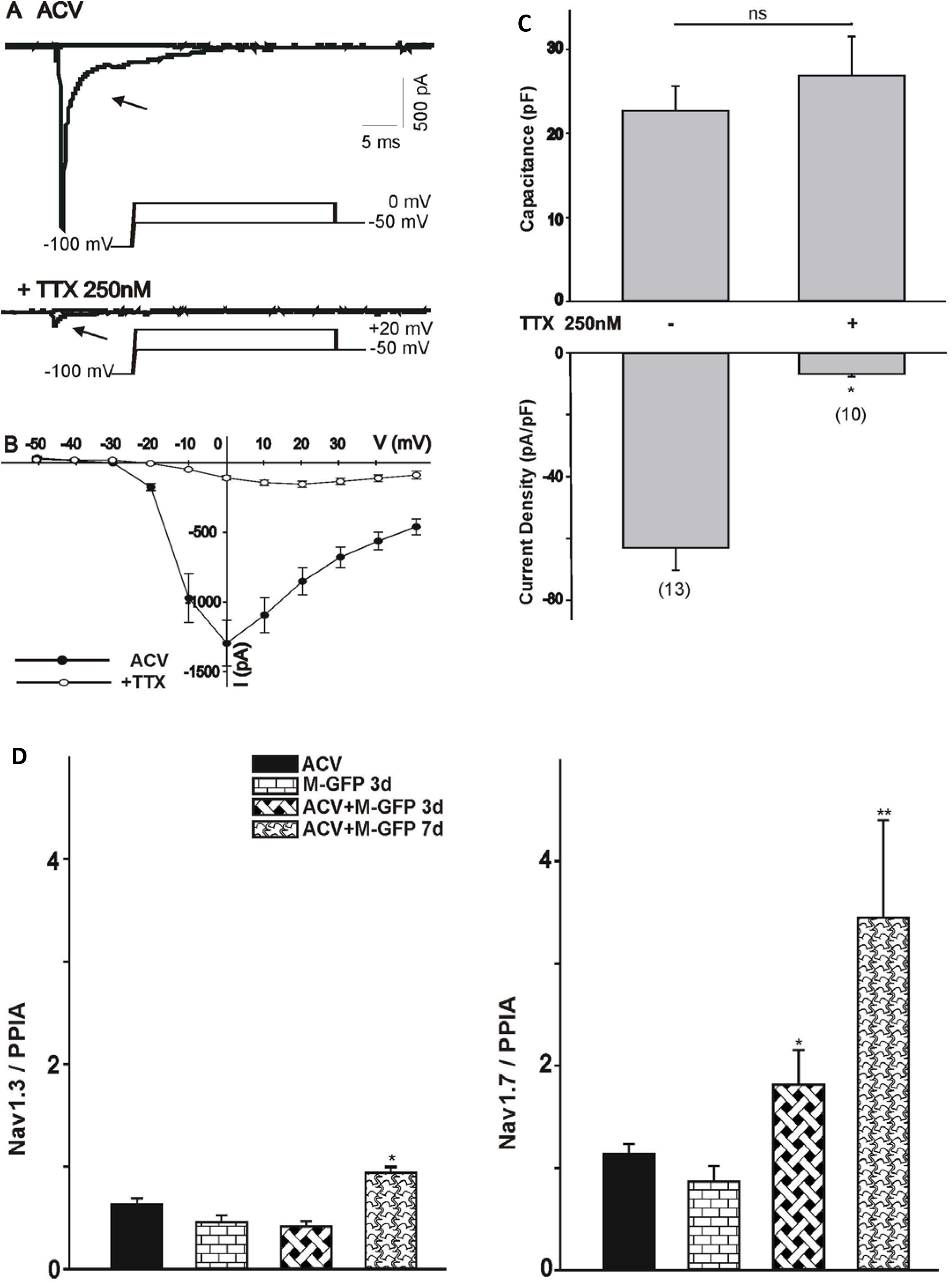
Subtype identification of voltage-gated Na^+^ channels in HD10.6 cells during HSV-1 infection. HD10.6 cells were cultured and tested in different conditions. A) The current traces of ACV group generated by voltage step to 0 mV from a holding potential of −100 mV was eliminated following TTX treatment. The peak current were obtained by voltage step to +20mV after 250nM TTX treatment; B) The current-voltage (I–V) relationship for the peak Na^+^ current amplitude in differentiated HD10.6 cells before and after TTX treatment. It was generated by a series of voltage steps from a holding potential of −100 mV. C) Capacitance and Na^+^ current densities were compared between cells with or without TTX treatment. The asterisk (*) denotes p < 0.05 vs. ACV control. NS indicates the two groups are not significant statistically. D) Nav1.3 and Nav1.7 expression was detected by qRTPCR during latency establishment. The asterisk (*) denotes p < 0.05 vs. M-GFP 3d. The double asterisk (**) denotes p < 0.05 vs. ACV+M-GFP 3d

### Nav1.7 expression was increased in latently infected neurons

We continued to study Nav1.7 based on the observation that HD10.6 cells displayed TTX-sensitive VGSC activity. The expression profile of sodium channels in the latent neurons was analyzed by Western blot using antibodies against the TTX-sensitive channel subunits Nav 1.7. The results showed that Nav1.7 protein was expressed in control cells and higher in latent neurons (Fig. 6A, lane 1 and 2). TSA slightly decrease the protein expression compared to control cells (Fig. 6A, Lane 3) but caused a significant reduction in cells with latent reservoir (Fig. 6A, Lane 4). Additional studies by qRTPCR revealed that the Nav1.7 transcript of latent cells displayed a 4.25-fold increase in comparison to others (Fig. 6B, Lane 2). These results suggested that Nav1.7 protein expression was elevated in latent neurons, probably due to increased transcription influenced by latent virus. Nav1.7 expression on individual neurons was further assessed by immunofluorescent microscopy and it showed that Nav1.7 expression in the latent neurons was as high as the uninfected ones if not increased (Fig. 6C). The signal, nonetheless, became weaker in neurons with viral reactivation triggered by TSA (Fig. 6D), suggesting viral replication may have effects to reduce the Nav1.7 expression.

**Fig. 6.**
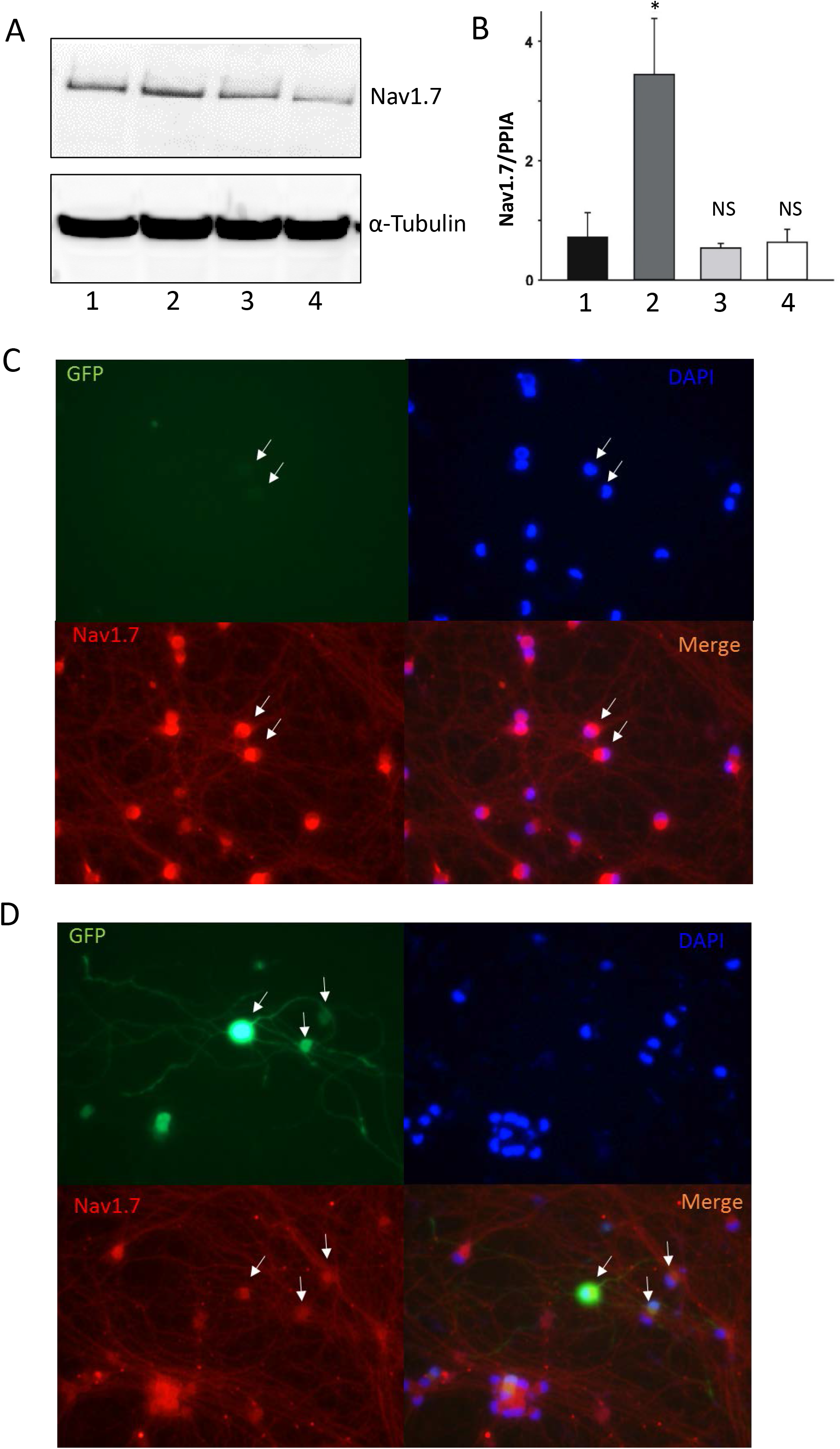
Expression of Nav1.7 in the HD10.6 cells. The expression of Nav1.7 protein and mRNA was assessed by A) Western blot and B) qRTPCR, respectively. Lane 1: Control; Lane 2: Latent cells; Lane 3: Control + TSA; Lane 4: Latent cells + TSA. The Nav1.7 expression was further evaluated individually by Immuno-fluorescent microscopy. C) Latent cells; D) Latent cells + TSA. The asterisk (*) denotes p < 0.05 vs. control. NS indicates there are no significant differences statistically comparing to control

### Latent neurons exhibited very high VGSC activity

Electrophysiological studies indicated that capacitance of cells under treatment did not change and Na^+^ current density from the latent neurons was increased approximately 20% in comparison to control cells (compare Fig. 7A and B). TSA treatment, nevertheless, decreased the current density by 50% regardless of the status of latency (Fig. 7B). Current density plot analyses studying individual neurons demonstrated that control cells (Fig. 7C) displayed a wide range of current densities between very high (current density > 80 pA/pF, 40%), high (current density between 60-80 pA/pF, 40%), and moderate (current density between 40-60 pA/pF, 20%) densities. Latent neurons conversely presented a unique pattern with approximately 90% of the neurons exhibiting very high current density (Fig. 7D). Note that for latent neurons, there were less than 0.02% of the cells with bright GFP, which were regarded as failing to establish latency, none of our current recordings are from those bright GFP cells. As for the TSA treatment, both control (Fig. 7E) and infected groups (Fig. 7F) revealed charts with current density shifting toward to the left with 45-50% neurons exhibiting low (<40 pA/pF) current densities, suggesting a trend of decrease VGSC number and activity. Collectively these results indicated that latent neurons, compared to controls, appeared to have higher VGSC activity, presumably resulting from increased Nav1.7 protein expression due to increased transcription influenced by latent virus. This Nav1.7 increase was nullified when reactivation occurred by TSA. However, it is important to note that TSA treatment *per se*, though did not considerably affect the level of protein expression, significantly decreased the VGSC activity as well.

**Fig. 7.**
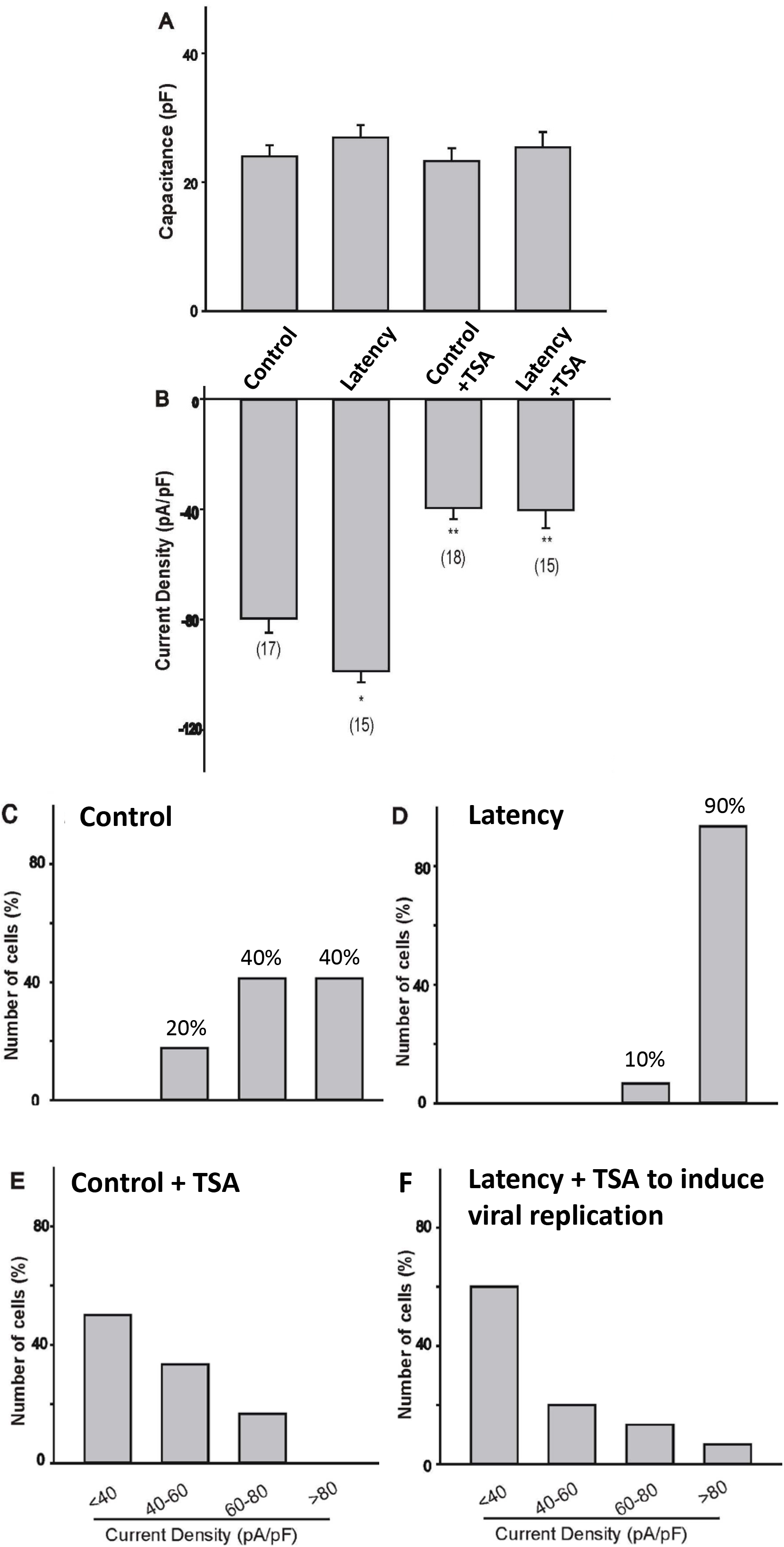
Evaluation of VGSC activity during latency and after induced viral replication. Na^+^ currents of the whole cell were recorded to measure the activity. A) Cell capacitance was measured and compared to controls. B) Na^+^ current density was calculated based on the peak current and capacitance. Panel C to F are current density plot analyzed to compare the VGSC activity of neurons between each group. C) Control; D) Latency; E) Control+ TSA; F) Latent cells + TSA. The asterisk (*) denotes p < 0.05 vs. the control. The double-asterisk (**) signifies p < 0.05 vs. the latent group

## Discussion

The glossary of viral latency and reactivation has been an interesting topic for all virologists especially the ones who studied alpha herpesviruses (29). It is generally believed that HSV-1, for example, exercised two alternative infection plans, lytic and latent infections. The episodes were launched by initial acute lytic infection marked by active viral replication in epithelial cells followed by latency established inside the sensory neurons of the ganglia, such as TG or DRG, innervating mucosa of the primary infection site (30). The operational definition of latency was suggested to be that a viral genome stayed and endured within a cell population in a time maintaining a dormant state without detectable lytic viral transcript, proteins, or infectious particles yet retained the capacity to replicate and reactivate (30–36). The best example recapitulating these features of HSV latency would be the *in vivo* rodent models (37–39). Besides the animal models, a number of cell culture models generated in neuroblastoma cell lines, differentiated embryonal carcinoma cell line, induced pluripotent stem cells, neuronal stem cells, and embryonic stem cells have been developed (35, 39–44). Although they reluctantly satisfied all characteristics of latency, these *in vitro* models offered a number of benefits in studying HSV latency particularly in understanding the molecular mechanisms.

The current cell culture model derived from human DRG was first used to study nociceptive properties (28). Recent progress through infecting with HSV-1 in the presence of ACV demonstrated a unique platform revealing many properties mimicking HSV-1 latency, such as silent phase of infection and low frequency reactivation (34). Our present results using strain McKrae largely agreed with this previous finding with additional observations. First, a quiescent infection was achieved without infectious virus detected inside or outside the cells but TSA treatment appeared to switch the latent reservoir into a replication mode, imitating the viral reactivation observed *in vivo*. Second, TSA action results in increased viral transcription and production of viral progeny within the cells, however, it fails to trigger the release of infectious viral particles. Last, the transcription of HSV-1 ICP0 and TK progressively diminished during the latency establishing stage. A slightly different pattern was observed in HSV-1 LAT, which temporarily accumulated at 3dpi followed by a steady decline thereafter. Noted that the PCR oligo primers were designed to target the LAT stable intron (45), it is not impossible that this accumulation resulted from delayed degradation. However, we cannot rule out the possibility that HSV-1 LAT may play a role in latency initiation, which warrant future studies.

Our results indicated that VGSC activity in HD10.6 cells was quickly eliminated by HSV-1 lytic infection, in consistent to the preclinical investigation that peripheral nerve injury decreased pain-associated voltage-gated channels in the injured DRG (12). Additional studies indicated that sodium currents gradually recuperated during the establishment of latency. Furthermore, the latent neurons demonstrated a unique functional expression profile in which 90% of the cells exhibited increased sodium currents with very high density of VGSC. This observation suggested that HSV-1 employed a strategy to facilitate the replication during lytic infection by bringing down VGSC activity in order to decrease neuronal excitability, then to sustain the quiescent state by promoting the functional expression of VGSC thus increased the excitability. In another word, increased neuronal excitability seemed to impede viral replication and participate in the maintenance of latency. It is further supported by previous report (18) and our observation that neuronal depolarization by potassium chloride significantly inhibits viral gene expression and replication (unpublished data). On the other hand, the sensation of neuropathic pain may be a part of innate immunity to suppress viral replication and held the virus in check of not actively replicating. The investigation is underway to understand the underlying molecular mechanisms.

The observation of Na^+^ current recovery in neurons following the establishing of latency prompted us to study which sodium channel subtypes may contribute to the rescue. The first examined subunit was Nav1.3 since it was reported to be upregulated in DRG neuronal injury and the protein was accrued in patients suffering painful neuropathy (12, 46–49). However, results examining transcript (by qRTPCR) and protein (by western blot and immunofluorescent microscopy) did not show any Nav1.3 expression (data not shown). Our investigation on Nav 1.7 indicated that its transcription was active and the total protein was slightly higher during latency but reduced upon TSA treatment, which causes viral reactivation. Immunofluorescent microscopy revealed that latent neurons under the influence of TSA displayed reduced expression of Nav1.7, suggesting that neurons with latent HSV-1 exhibited an induction of Nav1.7 and reactivation impeded this increase. Subsequent electrophysiological studies investigating the VGSC activity demonstrated that sodium current, though higher in latent neurons and lower after reactivation, it also revealed a decrease in the TSA control. Given the fact that total Nav1.7 protein was not significantly decreased by TSA, it is likely that TSA modulated cell signaling and obstructed the trafficking of Nav1.7 to the cell surface. TSA decreases the neuronal excitability and induce the viral replication, which indicated that decreased neuronal excitability may facilitate viral reactivation.

Nav1.7 has been shown to be physiologically associated with the function of other VGSC to amplify generator potentials and boost subthreshold stimuli thus causing VGSC activation, efficient recovery from inactivation, and generates high frequency action potentials (50). Blockade of the channel indicated that Nav1.7 participated in the key TTX-sensitive Na^+^ current in small-diameter DRG neurons (51). Nav1.7 activity has been linked to post-translational modification such as phosphorylation (52). Signaling transduction studies suggested that phosphorylation on an intracellular loop by the extracellular signal-related kinase1/2 (ERK1/2) influenced Nav1.7 activity, thus increasing its efficacy during action potential firing (52, 53). Other reports indicated that Fyn tyrosine kinase can phosphorylate Nav1.7, hence influenced channel expression and gating (54). It is unknown how HSV-1 latency can influence VGSC activity. Understanding the molecular mechanisms involved in the regulation of Nav1.7 activity may have important implication during the HSV-1 latency and reactivation.

Our current study demonstrated that differentiated HD10.6 cells displayed robust VGSC activity and steady recovery of Na^+^ currents after elimination by HSV-1 infection. Differentiated HD10.6 cells supported the establishment of a quiescent state of infection mimicking latency and maintained the capacity to replicate similar to reactivation induced by TSA. During the maintenance stage of dormant infection, the neurons harboring latent reservoir of virus appeared to have very high functional expression of VGSC, presumably to be Nav1.7, and reactivation decreased the VGSC activity. This abnormal pattern of Na^+^ channel expression profile following HSV-1 lytic infection, latency, and reactivation may disturb the ability of sensory neurons to transmit pain information. HSV-1 infection often alter pain sensory transmission with both diminished and enhanced pain sensation (8, 16, 55). Functionally, elevated sensory neurotransmission and increased excitability due to higher expression of voltage-activated Na^+^ currents during latency could potentially enhance pain signaling but offer a stealth capacity escaping detection by the host immune system. This unique change in ion channel expression at different stages of the viral infection as a result of host responses to virus assault remains to be investigated and may have implication for the development of therapeutic protocols to treat herpes virus-mediated pain, trigeminal neuralgia, and post herpetic neuralgia.

## Materials and Methods

### Cells and virus

The human DRG neuronal cell line HD10.6 was used as an in vitro model as previously described (33, 34) and a gift from Celgene Corporation (San Diego, CA). African green monkey kidney Vero cells (ATCC, Cat# CCL-81) and the culture condition was described previously (56, 57). Only cells passaged less than 15 times were used in this work. None of the cell lines used in this work has been misidentified according to the International Cell Line Authentication Committee (ICLAC) and cell line authentication was performed by the providers using short-tandem repeat (STR) analysis. HSV-1 strain McKrae virus (M-GFP) with the GFP expression under the control of a cytomegalovirus (CMV) promoter was used throughout the current study (58).

### Infection, latency establishment, and reactivation

For HSV-1 lytic infection, differentiated HD10.6 cells (differentiation ≥7 days) were exposed to HSV-1 for 1 h at a multiplicity of infection (MOI) of 1. Unbound viral particles were washed out after 1 h and fresh differentiation media was applied followed by culture for another 3d before used in subsequent experiments. To establish HSV-1-latently infected cultures, differentiated HD10.6 cells were pre-treated with 100 μM Acycloguanosine (acyclovir or ACV, Cat#: A4669, Sigma-Aldrich) overnight, then exposed to HSV-1 for 1 h at a MOI of 1. Unbound viral particles were washed out after 1 h and differentiation media (DM) supplemented with 100 μM ACV was applied for 3d, then changed with fresh DM with 100 μM ACV followed by culture for another 4d to allow for the establishment of HSV-1 latent infection. ACV was then removed by vigorous wash with DM to allow for the maintenance of HSV-1 latent infection, which was defined as extremely weak or no GFP expression and low viral gene expression with no infectious HSV-1 viral particles release for a prolonged period of time. The Reactivation of HSV-1 from latency was tested by adding 1μM TSA (Cat#: T1952, Sigma-Aldrich) or 20μM LYS294002 (Cat#: 440202, Sigma-Aldrich) for 2d.

### Fluorescent microscopy

Morphometric analysis and GFP fluorescence of cultured HD10.6 cells were obtained with a Nikon Eclipse Ti microscope with a 20x inverted objective and a Photometrics Coolsnap EZ cooled camera. The intensity of GFP fluorescence change indicate the change of HSV-1 activity in infected HD10.6 cells and the protocol was previously described (26, 59).

### Quantitative PCR

Quantification of viral gene expression was performed as previously described (26, 45, 57). Briefly, RNA was collected using iScript sample preparation reagent (Cat#: 170-8898, BioRad) followed by reverse transcription using iScript RT Supermix (Cat#: 17008841, Bio-Rad) to produce cDNA template. The qPCR reactions were performed using the SsoAdvanced Universal SYBR green supermix (Cat#: 1725271, Bio-Rad) on triplicate samples using primers specific for the selected viral or human genes listed in Table 1. The qPCR reactions were carried out at 95°C for 30s, 95°C for 10s, followed by annealing temperature for 30s (39 cycles), annealing temperature for 5s. All of the different gene mRNA expression was normalized by peptidylprolyl isomerase A (PPIA) expression.

**Table 1:**
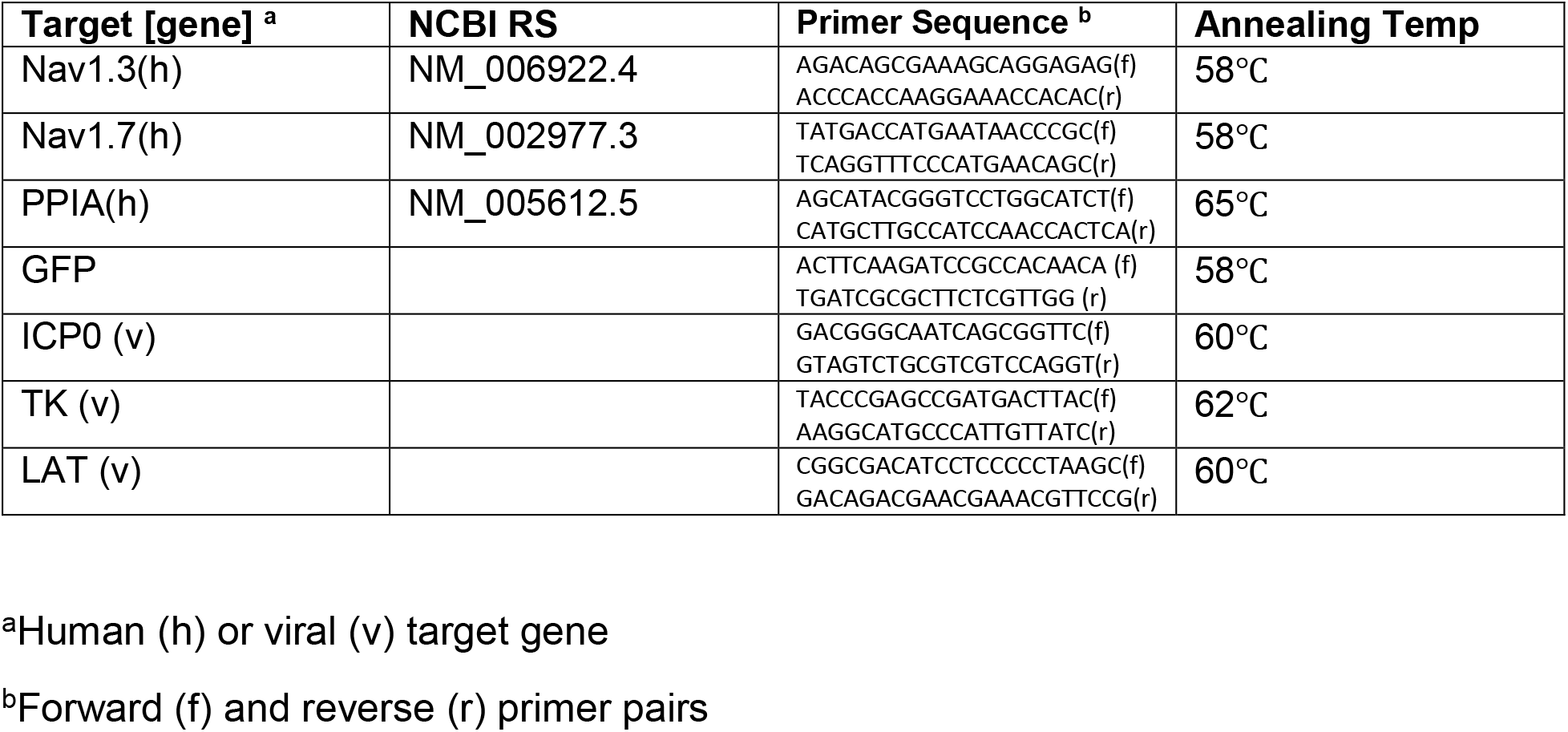

### Plaque assay

The plaque assays were performed as previously described (26, 56, 57). Briefly, the cell lysates of infected HD10.6 cells were mixed with 500μl media and subjected to three cycles of freeze-thaw to have virus released into the media. The cell lysates as well as the supernatant from infected cultures were serially diluted before being added to Vero cell monolayer in 24-well plates seeded at 8×10^4^ cells/well. After 1hpi, the media were removed and replaced by fresh culture media supplemented with 5% FBS. The cells were fixed after 2d with ice-cold methanol and stained with crystal violet prior to counting plaque formation. Data was collected from triplicates.

### Electrophysiology

Visualization of HD10.6 cells during recordings was performed by A Nikon Eclipse Ti inverted microscope (Nashua, NH) equipped with Hoffman optics and epifluorescence filters, which can be used to identify the infected HD10.6 cells by the expression of GFP. Voltage-activated sodium currents from individual cells was recorded using the whole-cell patch-clamp technique. Recordings were performed at room temperature (22-24°C). The pipette electrode solution consisted of (in mM) CsCl (120), MgCl_2_ (2), HEPES-KOH (10), EGTA (10), ATP (1), and GTP (0.1), pH 7.4 adjusted with CsOH. The extracellular solution used to measure Na^+^ currents contained (in mM) NaCl (145), KCl (5.3), CaCl_2_ (0.54), MgCl_2_ (5.7), HEPES (13) and Glucose (5), pH 7.4 adjusted with KOH. Na^+^ currents were generated by applying a 20 ms-depolarizing step to various potentials from a holding potential of −100 mV. A MULTICLAMP 700A amplifier was used to apply voltage commands and PCLAMP software (Axon Instruments, Foster City, CA) was used to perform data acquisition and analysis. A MultiClamp 700B Commander was used to compensate pipette offset, whole cell capacitance, and series resistance (≤10 MΩ) automatically. Sampling rates were 5~10 kHz. For quantitative analysis, cell size was normalized as previous described (60, 61). In short, dividing the recorded current amplitudes by cell capacitance, which was determined by integration of the transient current evoked by a 10-mV voltage step from a holding potential of −60 mV. Voltage-gated Na^+^ current density was calculated by dividing the maximal current amplitude generated by a voltage step around +10mV from a holding potential of −100 mV by the cell capacitance. TTX was purchased from TOCRIS (Cat#: 1069).

### Western blot analysis

Immunoblot analysis of the voltage-gated Na^+^ channel subunit Nav1.7 was conducted as previously described (26), using a rabbit anti-Nav1.7 primary antibody (1:1000, Cat#. ab196906, Abcam, US). Proteins were separated on pre-cast SDS-PAGE 8% gels (Cat#: NW0080, ThermoFisher) and then transferred to nitrocellulose membranes using the Invitrogen iBlot system. After a horseradish peroxidase-conjugated anti-rabbit secondary antibody (Cat.#211-035-109, Jackson ImmunoResearch Lab, West Grove, PA) incubation for 1 h at room temperature, a chemiluminescent substrate (Super Signal West Pico, Cat.#34080, ThermoFisher) was applied, the signals were captured using a ChemiDoc™ XRS+ documentation system (Bio-Rad, Hercules, CA). For controls, membranes were stripped and reprobed with a tubulin-specific antibody (1:2000, Cat#: 05-829 Millipore), followed by incubation with a peroxidase-conjugated anti-mouse secondary antibody (Cat#115-035-146, Jackson ImmunoResearch Lab, West Grove, PA) and chemiluminescent detection. Densitometry analysis was used to determine the changes in Nav1.7 protein expression using Image Lab software (Bio-Rad).

### Immunofluorescent Assays

The immunofluorescent assay was performed as previous decried by Figliozzi et al. (2014). Briefly, infection of HD10.6 cells seeded on Matrigel and Poly-D-Lysine pre-coated 24 well-plates was performed before fixation of cell cultures. Cells were washed with PBS 2X 5 mins before applying 4% paraformaldehyde (PFA) to fix the cells. After 15 mins incubation at room temperature, cells were washed with PBS 3X 5 mins. Permeablize the cells with 0.25% triton X-100 (diluted in PBS) for 5 mins and wash with PBS for another 3X 5 mins. Then the cells were incubated with blocking buffer (1XPBS, 1% BSA, 0.3M glycine, 0.1% Tween 20) supplemented with 5% goat serum for 1 h at room temperature. After overnight incubation with Nav1.7 antibody (Cat#: ASC-008, Alomone, Jerusalem, Israel, 1:250, diluted in antibody dilution buffer: 1X PBS, 1% BSA, 0.1% Tween 20) at 4°C, wash with PBS 3× 5 mins. Followed by incubation with goat anti-rabbit lgG H&L (Alexa Fluor® 594) preadsorbed secondary antibody (1:500 diluted in antibody dilution buffer, Cat #: ab150088, Abcam) for 2 hrs at room temperature in dark. After three times wash with PBS, cells were counterstained with 4’-6-diamidino-2-phenylindole (DAPI) (1μg/ml in PBS) for 10 minutes in dark. After three times wash with PBS, the plate was ready for imaging. Images were obtained with an Olympus IX2-RFACA motorized fluorescence microscope with a 40 x inverted objective. All of the images were analyzed and colorized using Nikon NIS-Elements imaging software.

### Data Analysis

Statistical analyses consisted of Student’s unpaired t-test and one-way ANOVA followed by post hoc analysis using Tukey’s honest significant difference test (STATISTICA software, Tulsa, OK). All data values are presented as mean ± SEM, p ≤ 0.05 was regarded as significant.

## Acknowledgement

We are grateful for the gift of HD10.6 cells from the Celgene Corporation (San Diego, CA). We thank Catherine Pearce and Nicholas Baird from the University of Colorado, School of Medicine for the scientific support. This project was supported by the University of Maryland Eastern Shore School of Pharmacy and NINDS/NIH grant R01NS081109 to SVH. The content of this work does not necessarily represent the official views of the NIH and is solely the responsibility of the authors. NINDS had no role in study design, data collection and analysis, decision to publish, or preparation of the manuscript.

## Conflict of Interest

The authors declare no conflict of interest.

## Author contributions

SVH started the project with the initial hypotheses and funding acquisition. MMC and SVH designed the experimental approaches further modified by QZ and FC. All cells and viruses were cultured, differentiated, and maintained by QZ and FC. Infection, gene expression analyses, and plaque assays were performed by QZ, FC, and SVH. Electrophysiology was performed by QZ with assistance by MMC. Immunofluorescent assays were performed by QZ and SVH. The results were interpreted and discussed by QZ, FC, MMC, and SVH. The manuscript was written by QZ and SVH, with help and discussion by FC and MMC.

